# Stretchable Composite Acoustic Transducer for Wearable Monitoring of Vital Signs

**DOI:** 10.1101/2019.12.14.876409

**Authors:** Yasin Cotur, Michael Kasimatis, Matti Kaisti, Selin Olenik, Charis Georgiou, Firat Güder

## Abstract

We report a highly flexible, stretchable, and mechanically robust low-cost soft composite consisting of silicone polymers and water (or hydrogels). When combined with conventional acoustic transducers, the materials reported enable high performance real-time monitoring of heart and respiratory patterns over layers of clothing (or furry skin of animals) without the need for direct contact with the skin. Our approach enables an entirely new method of fabrication that involves encapsulation of water and hydrogels with silicones and exploits the ability of sound waves to travel through the body. The system proposed outperforms commercial, metal-based stethoscopes for the auscultation of the heart when worn over clothing and is less susceptible to motion artefacts. We have tested the system both with human and furry animal subjects (*i.e.* dogs), primarily focusing on monitoring the heart, however, we also present initial results on monitoring breathing. Our work is especially important because it is the first demonstration of a stretchable sensor that is suitable for use with furry animals and do not require shaving of the animal for data acquisition.

## 1. Introduction

In modern global healthcare, both for humans and animals, wearable devices are expected to play a major role for remote and continuous monitoring of health and the early detection of diseases^[1]^ The increasing cost of healthcare, limited access to health centers (*e.g.*, due to long waiting times), and aging populations are the main drivers of wearable health monitor development.^[2]^ The recent advances in wearable monitoring tools, however, were only made possible with the emergence of low-cost mobile devices (*e.g.*, smartphones).^[3]^ When combined with mobile devices, wearable health monitors can acquire, manipulate, store and transmit data inexpensively compared to conventional solutions.^[4]^ At the same time, flexible and stretchable devices could provide unobtrusive and continuous monitoring of health without sacrificing comfort.^[5–12]^

(Opto)electronic sensors integrated into flexible/stretchable materials,^[13,14]^ with a wearable form factor, offer an accurate yet inexpensive method to monitor vital signs of health (heart rate in particular) non-invasively.^[15]^ The biggest shortcoming concerning these sensors (such as wearable optoelectronic heart rate monitors), however, is that they must be worn over the bare-skin of the user for reliable collection of data. Users may even need to shave parts of their body to be able to make accurate measurements.^[16]^ This produces inconvenience, discomfort, and often causes users to stop wearing their devices (since they continuously need to shave). Photoplethysmography (PPG) and electrocardiography (ECG) are two popular methods used in wearables for measuring heart rate (HR). While PPG is an optical method that detects HR by way of detecting the rate of blood flow, ECG captures electrical signals generated by the contraction and relaxation of the heart muscles.^[17],[18]^ Both methods generally require hair-free skin; often, compensation for motion artefacts is also needed. Most wearables for measuring heart rate require direct contact with the skin. Materials with low compliance do not conform to the curvature of the body; therefore, this produces poor-quality or faulty data, the devices are less comfortable and may require additional materials (such as conductive gels) for the measurement.^[19]^ There are also remote methods of measuring HR such as ballistocardiography (BCG) and seismocardiography (SCG) which provide unobtrusive collection of data but they fail to function when subjects move, due to motion artefacts.^[20]^

The use of stretchable materials with high compliance for the fabrication of wearable devices provides the means for creating robust yet comfortable tools for continuous monitoring of vital signs such as heart rate.^[21]^ Conventional electronics can be encapsulated with soft materials to improve conformation to the curvature of the skin and may be worn like a tattoo.^[22]^ Conformal and direct contact with the skin using wearable tattoos also enables alternative modes of monitoring heart rate including acoustic or other pressure-based sensing modalities.^[23],[24]^ These devices, however, still require *direct* (hard) contact with the skin and generally fail when the tissue is wet; they may also cause irritation/allergic reactions. Furthermore, sophisticated “tattoo-like” stretchable devices require advanced methods of microfabrication (hence expensive) and are meant to be thrown away after a single use (*i.e.* disposable).

In this article, we report a stretchable wearable composite acoustic transducer for monitoring vital signs. The composite device uses water or hydrogels as the medium for the propagation of acoustic waves which are encapsulated inside a stretchable silicone membrane (note that water/hydrogels are not applied directly to the skin). The acoustic signal originating from within the body are captured using battery-operated conventional microelectronics that can monitor HR and respiration over long periods, store, analyze and transmit the data to a nearby computer wirelessly. Unlike most wearable health monitors, our system can be worn over clothing or hairy skin (coat/fur) of animals such as working, military or police dogs without the need for shaving the animal. In this work, we particularly focus on monitoring HR (and only provide preliminary results for monitoring respiration) and characterize the system developed for both dog and human subjects.

## 2. Results and Discussion

### 2.1. Fabrication of the transducer

Figure 1 illustrates the four-step fabrication procedure for the water-silicone composite acoustic transducer. In this scheme, (i) a polylactic acid (PLA)-based polymer mold is 3D printed to cast the bottom-outer layer of the silicone membrane (2 mm in thickness). (ii) The silicone membrane produced is filled with deionized water (or hydrogels) and, (iii) uncured silicone is poured directly on top of the water to fully encapsulate it (see supporting information – SI – **Video SI-V1** for the production of water-silicone composite transducer). During this process, the silicone introduced naturally spreads itself over the water added inside the bottom-outer silicone membrane (*i.e.* does not mix) and forms another thinner membrane (20-100 µm) which covers the entire top-surface and is highly stretchable and flexible (**Figure S1**). This process, though may appear counter-intuitive, works extremely effectively and allows the silicone membrane from curing and bonding with the top edges of the bottom layer. This method also works if the deionized water is replaced with a hydrogel (**Figure S2**), producing a hydrogel-silicone composite. Once cured, the resulting structure contains no air bubbles (air bubbles reduce the quality of data acquired acoustically). (iv) In the final step, a conventional microphone with its associated electronics is placed on the outside of the silicone membrane and encapsulated with an additional layer of silicone to produce a monolithic device (**Figure S3**). To create a wearable form-factor, the water-silicone transducer can also be monolithically integrated into a silicone-based stretchable harness (Figure 1) which contains additional custom-built, battery-operated wireless electronics (**Figure S3**). The final system produced can be worn over the chest and conforms to the curvature of the body, allowing acquisition of acoustic signals from the subject.

**Figure 1.**
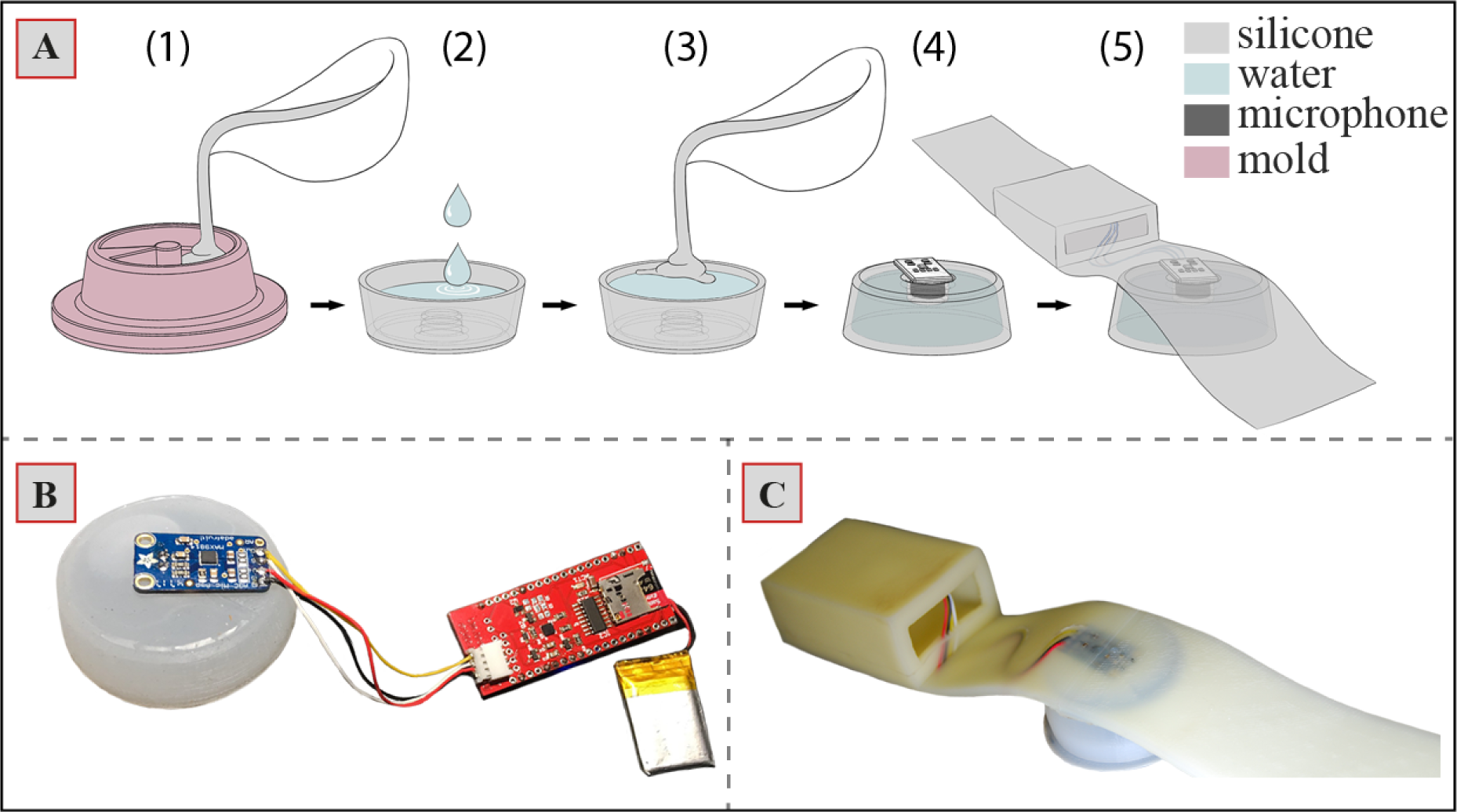
**(A)** Fabrication steps of the water-silicone composite transducer: 1) Degassed, uncured liquid silicone is poured in the mold and left to cure partially for two hours. 2) Partially cured silicone is removed from the mold and filled with water. 3) Uncured, liquid silicone is poured on the water; silicone spreads itself over the water and continues to cure with the partially-cured part, fully encapsulating the water. 4-5) Microphone is placed in the recess and buried in more silicone to create a monolithically wearable harness; **(B)** Photograph of the wireless electronics, battery and microphone; **(C)** Photograph of the harness produced by embedding the microphone amplifier with silicone. Electronics and battery are placed in a 3D printer container and placed in the sleeve on the harness.

### 2.2. Characterization of performance

We investigated the performance of the composite acoustic transducers for monitoring heart sounds, *i.e.* phonocardiography (PCG). PCG is a powerful method that produces a graphical recording of all sounds and murmurs originating from the flow of blood and motion of valves in the heart.^[25]^ There are two major heart sounds: (i) the lub sound – S1 – arises from the closure of mitral and tricuspid valves in the beginning of systole; (ii) the dub sound – S2 – occurs during the closure of aortic and pulmonic valves.^[26]^ A successful PCG device should be able to resolve these sounds accurately with little noise (most PCG devices are susceptible to movement) which can be used for measuring HR or making diagnosis of diseases of the heart.^[27–29]^

In the first experiment, we characterized the composite transducers using simulated heart sounds (see **Figure S4** for the experimental setup). To benchmark the quality of the sounds recorded with composite transducers and understand the effects of materials/geometry on transduction, we produced seven different types of silicone composite transducers (see Figure 2A for time series data) using the following materials: (i) commercial, non-flexible and non-stretchable stethoscope diaphragm, (ii) air-silicone composite with 15 mm height – this material is similar to the water-silicone composite but instead is not filled with water and contains air, (iii) all-silicone transducer with 15 mm height – consists of a block of silicone, (iv) water-silicone composite with 15 mm and (v) 30 mm height of water. We downloaded a reference PCG recording of a healthy heart beating at 60 beats/min from an online repository by ThinkLabs Medical LLC.^[30]^ Most heart sounds are observed between 20-100 Hz as shown in the frequency response of the original sound signal (see **Figure S5** for original heart sound). We played this sound using a loudspeaker and re-recorded the sounds generated using the transducers (see **Figures S6 to S9** for time series and **S10-S13** for the frequency spectra of the recordings).

**Figure 2.**
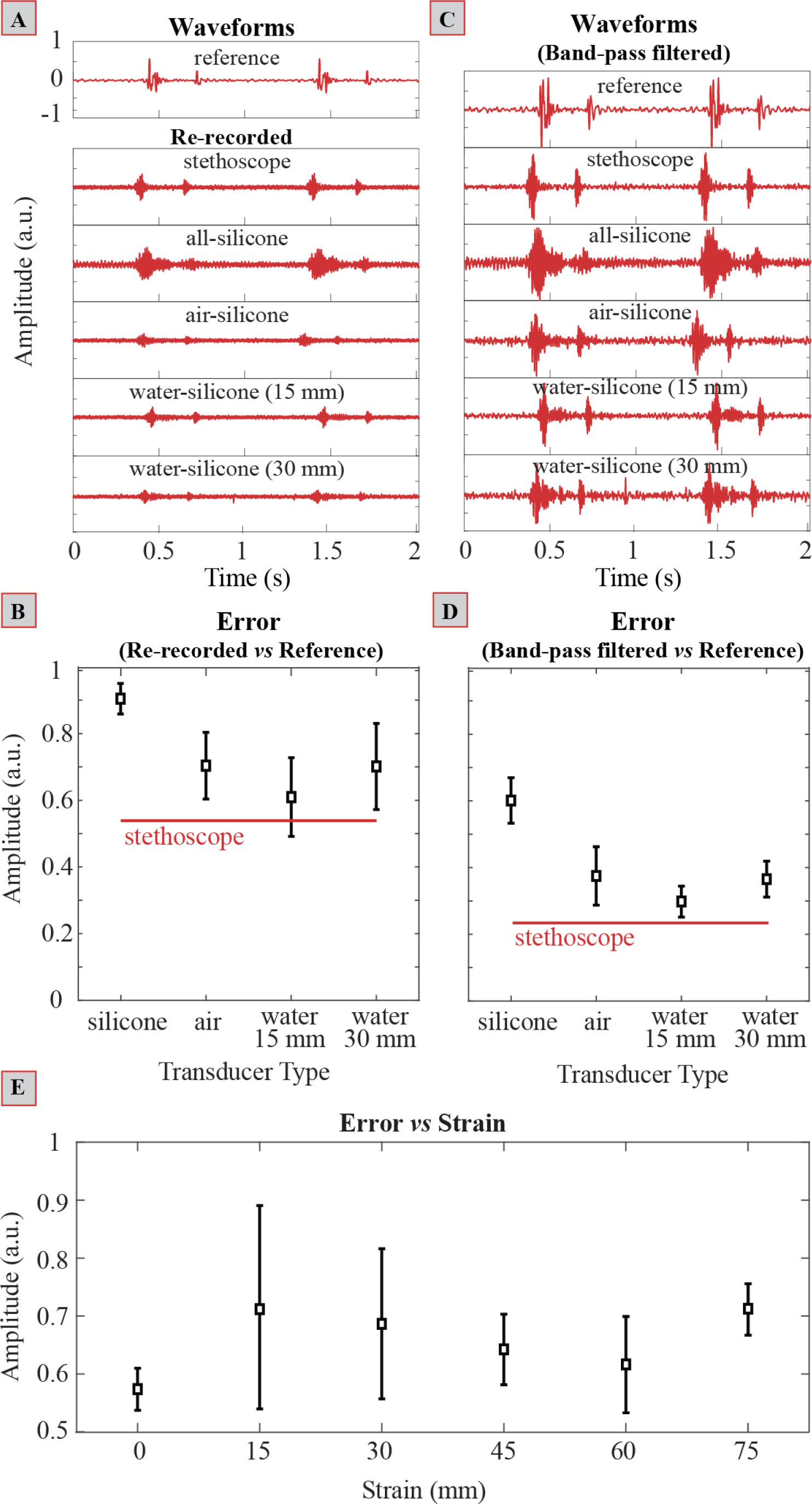
**(A)** Recording of simulated heart sounds. The simulated heart sounds (reference recording downloaded from an online repository) were played through loudspeakers and re-recorded using transducers made of different materials. Air-silicone and all-silicone transducers were both 15 mm in height whereas for water-silicone composite transducers, the heights were 15 mm and 30 mm; **(B)** Error between the re-recorded sounds, shown in (A), and reference recording, calculated using dynamic time warping (DTW); **(C)** Normalized and filtered waveforms from (A) – Passband: 20 – 100 Hz. **(D)** Error between the bandpass filtered, re-recorded normalized sounds, shown in (C), and bandpass filtered, normalized reference recording calculated using DTW. **(E)** Error between re-recorded sounds and original recording calculated using DTW as a function of strain.

Visual evaluation of the time series data alone indicates that the air-silicone composite and all-silicone transducers produce recordings with lower quality; the air-silicone transducer attenuates the signal of interest and S2 is barely visible due to the noise. The all-silicone transducer creates little attenuation but introduces significant noise to the recording, reducing overall quality. To produce a quantitative measure of similarity, we used <u>D</u>ynamic <u>T</u>ime <u>W</u>arping (DTW) to calculate the error between the reference recording that is played back, and the signal captured by the transducers. DTW (see **SI-D1** for more information on DTW) is widely used as a similarity measure especially for biological signals such as speech.^[31–33]^

The DTW analysis shown in Figure 2B indicates that the water-silicone composite transducers (n=7) produce the smallest mean error among all stretchable materials, in comparison to the reference recording. Although the all-silicone transducers have the lowest variability, they exhibited the worst overall performance. Water-silicone composite transducers with a height of 30 mm and air-silicone composite transducers produced signals with higher mean error in comparison to the 15 mm water-silicone composite transducer. The transducer with the commercial stethoscope diaphragm performed the best among all devices tested; this result was expected since the attenuation of acoustic waves is lowest in dense solids. Of course, the commercial device is not stretchable, does not conform to the human body, and is susceptible to motion artefacts (moves easily), thus it is not suitable for continuous wearable sensing.

The degree of attenuation that an acoustic wave experiences depends on two factors: (i) the acoustic impedance which is the product of the density of the medium of propagation and speed of sound in that medium^[34]^ (ii) the coupling between the mediums of propagation if the waves are traveling across an interface.^[35]^ Unstable contact between (solid) mediums also produces noise that reduces the quality of a signal. Although in terms of the acoustic impedance of the medium, the all-silicone transducer had an advantage over the other silicone-based devices because it is less flexible and less stretchable compared to the air-silicone and water-silicone composite transducers, the coupling between the mediums was poorer due to the reduced contact area. This resulted in significant error during transduction (see **Video SI-V2** for comparison of stretchability). The 15 mm water-silicone composite transducer performed the best, most possibly due to the following three reasons: (i) higher flexibility/stretchability due to the thin silicone membrane compared to all-silicone transducers, providing better coupling between the transducer and loudspeaker (ii) water provides a medium of propagation with a lower acoustic impedance compared to air,^[36]^ and (iii) increasing the height of the water-silicone composite transducer from 15 mm to 30 mm also increases the length of propagation the acoustic waves need to travel from the surface of the speaker to the microphone, reducing signal-to-noise-ratio. Since the silicone-water composites have a positive Poisson’s ratio, we did not reduce the thickness lower than 15 mm as this would bring the microphone too close to the surface when stretched which could induce noise.

### 2.3. Enhancement of transduction performance using digital filters

We applied a bandpass filter (infinite impulse response Chebyshev filter with a passband of 20 – 100 Hz; Figure 2C) to the as recorded signals. Through visual inspection of the time series data, it is obvious that after filtering, the signal acquired from the all-silicone transducer remained noisy. The amplitude of the signals recorded with the air-silicone composite transducer exhibited an increase in quality and the S1 and S2 could now be clearly identified. The quality of the recordings measured by the 15 mm water-silicone composite transducer resembled closely the original heart sound and the 30 mm composite had higher noise. Once again, using DTW we have quantitatively measured the degree of similarity between the original recording and the measurements made with the transducers produced (Figure 2D). Digital filtering improved the quality of all recordings: the 15 mm performed the best, similar to the commercial stethoscope diaphragm but with the added benefit of being stretchable/flexible which would provide improved coupling when worn over clothing or a patch of hairy skin. In the remaining experiments, therefore, we used the water-silicone composite transducer with 15 mm thickness.

### 2.4. Effect of strain

To understand the effect of stretching on the performance of the 15 mm water-silicone composite transducer, we placed the devices to be tested on a loudspeaker (Harman Kardon Onyx Studio 4) with a curved surface. We stretched seven samples of water-silicone composite transducers at six levels of strain up to 75 mm to measure error with DTW (Figure 2E). The DTW results show that stretching does not produce a significant change in the performance of the water-silicone composite transducers, although there is a slight increase in variability/error when the devices are stretched without a trend. Nonetheless, we can conclude that stretching has a negligible effect on performance.

### 2.5. Testing with healthy human volunteers and dogs

In the next experiment, we tested the 15 mm water-silicone composite transducer to record the heart sounds of healthy human volunteers (n=5) wearing their daily clothing. For example, the subject shown in Figure 3A was wearing three layers of clothing on the day of the experiment (see **Figure S14** for comparison of acquired signals versus layers of clothing). All recordings (see **Video SI-V3**) were made in a standard laboratory environment with ambient noise. The heart sounds recorded before and after digital filtering, while the subject was standing, are shown in Figure 3B. We included the waveforms recorded by a commercial stethoscope for comparison. Since the commercial stethoscope was rigid and did not conform to the body well, the amplitude of the signals recorded were low, and the heart sounds were difficult to detect. Even though we were able to recover the S1 sounds by bandpass filtering (Butterworth filter with a passband of 20 – 150 Hz), the S2 sounds were still not visible. In contrast, both the S1 and S2 sounds of the heart could easily be identified with the water-silicone composite transducer. With the stretchable composite transducer, the recordings had a much better signal-to-noise profile (**Figure S15**) most probably due to the improved contact (*i.e.* improved conformation to the shape of the body of the subject). We did not observe a significant change in the quality of the signal recorded with the water-silicone composite transducer before and after filtering. We also applied wavelet denoising and envelope detection algorithms to obtain heart rates from the waveforms recorded during sitting or standing (Figure 3C **and Figure S16)**. For further validation of our method we measured the electrical activity of the heart using conventional ECG (note that the ECG electrodes were attached directly on the skin) while making PCG recordings with the water-silicone composite transducer. As shown in Figure 3D, both recordings produced highly correlated signals as reflected by the heart rates determined during the measurements (60 bpm).

**Figure 3.**
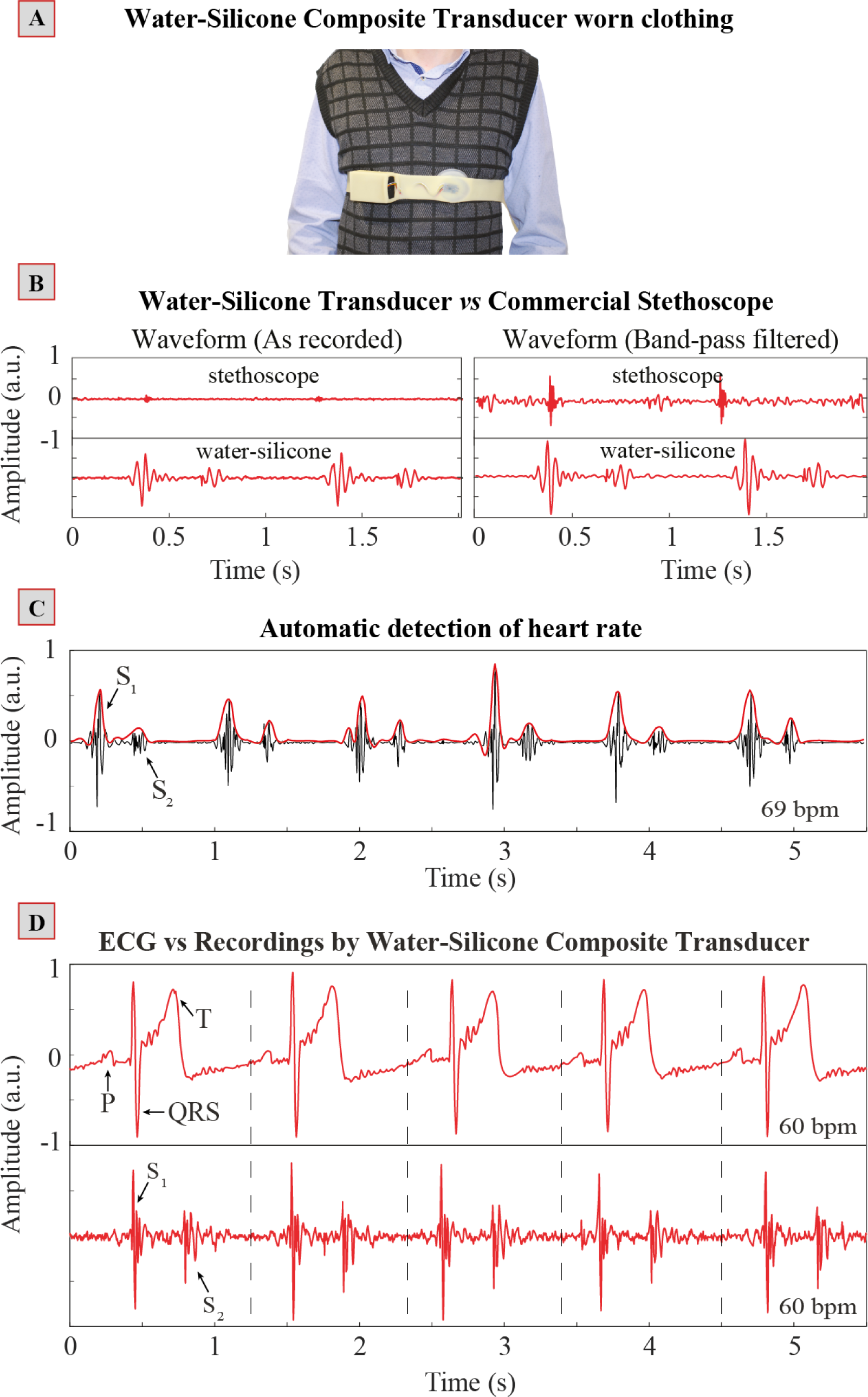
**(A)** Testing with healthy human volunteers. The water-silicone composite transducer is worn over layers of clothing; **(B)** As recorded sounds from the body with the water-silicone composite transducer versus commercial stethoscope – left. After bandpass filtering with a Butterworth filter (passband 20 – 150 Hz), the sound quality improves further – right; (**C)** Algorithmic detection of S1 and S2 waveforms recorded from a human subject and subsequent identification of heart rate. **(D)** Simultaneous recording of ECG (using commercial electrodes attached directly on the skin) and PCG (recorded with the water-silicone composite transducer) signals showing functional agreement.

In addition to that of human volunteers, we also tested the water-silicone composite transducer for monitoring the heart rate of a dog (Figure 4A). The test subject was a healthy Labrador Retriever. Unlike humans, most dogs have a thick coat of fur, rendering our method of measuring heart sounds particularly suitable for monitoring the heart rate of furry animals. Figure 4B shows the sounds of the heart of the dog which were bandpass filtered (passband: 20-150 Hz) to remove unwanted signals – *e.g.*, breathing sounds, ambient noise. Both the S1 and S2 regions can be clearly identified in the resulting waveform.

**Figure 4.**
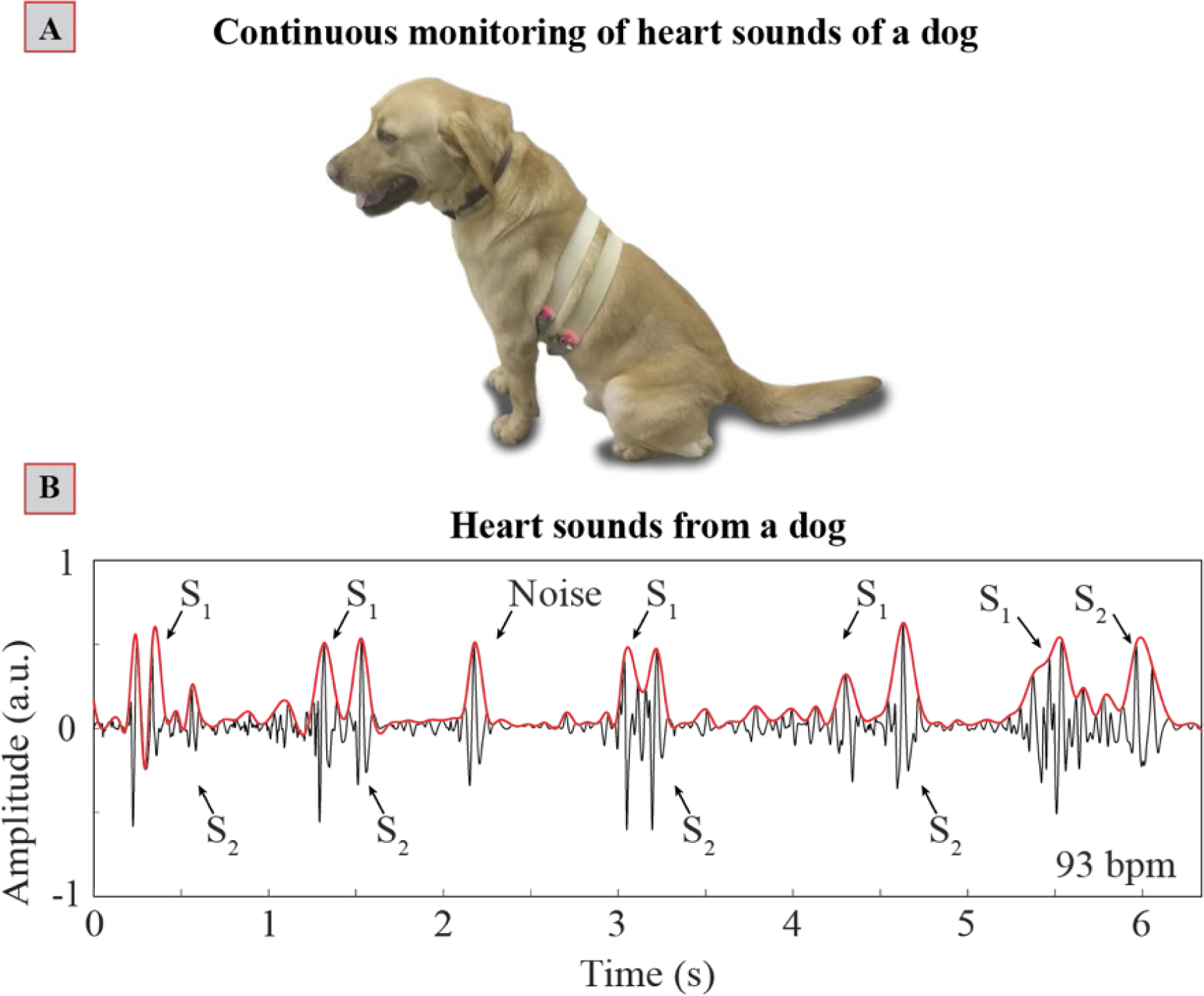
**(A)** Testing of water-silicone composite transducers worn over the furry coat of a dog; **(B)** Algorithmic detection of S1 and S2 waveforms recorded from the dog and subsequent identification of heart rate.

Overall, the wearable water-silicone composite transducers performed well with healthy human and animal subjects and allowed recordings of PCG continuously without the need for shaving or using conductive gels.

## 3. Conclusions

The water-silicone composite acoustic transducer is an unobtrusive, wearable measurement tool that can be used for monitoring vital signs continuously. Although in this work we have mainly focused on measuring heart sounds, as the spectral distribution of frequencies belonging to breathing and heart sounds/murmurs are different from each other, the breathing patterns can also be measured through the use of digital filters (**Figure S17)**.^[37]^

The system is sufficiently low-cost ($0.5 for silicone, $30 for the electronics, battery and microphone) compared to a commercial electronic stethoscope and easy to fabricate using widely accessible prototyping tools (*e.g.*, 3D printers). The use of an Internet-of-Everything system based on the inexpensive ESP32 board allows acquisition and transmission of recordings to a remote device wirelessly. The digital recordings can be pre-processed using the on-board microcontroller and further processed on a nearby (more powerful) mobile device with cloud storage capabilities. Machine learning algorithms can be applied to the data collected to detect anomalies and alert healthcare professionals/veterinarians (or pet owners) of illnesses.

The wearable system, in its current format has at least four disadvantages: (i) Silicone elastomers require curing (i.e. slow) and are not compatible with low-cost mass manufacturing methods such as injection molding. (ii) Acoustic transduction is susceptible to external environmental sounds which reduce the quality of the recordings although the contribution of the external sounds could be reduced through further soundproofing. (iii) When used with animals such as dogs the system has difficulty collecting high quality heart sounds from dogs that pant strongly. (iv) Sudden movement of animals (especially highly active dogs) may also introduce additional noise to the recordings through physical contact with external objects or body parts (such as legs). These disadvantages can, however, be largely eliminated by the use of additional sensors to measure breathing/panting (such as elastomeric strain sensors recently developed in our group)^[38]^ and a different form-factor for the harness.

The system proposed is suitable for use with humans and expected to work with a range of animals such as furry pets, working animals (such as sniffer dogs) or livestock although in this work, we only performed testing with a dog. In the future, the wearable composite device presented may provide an unobtrusive alternative to current health monitors for human and animal care at a lower cost.

## 4. Methods and Materials

### 4.1. Non-invasive testing with human volunteers and dog

Prior to non-invasive human and dog experiments, a detailed risk assessment was performed according to the guidelines provided by Imperial College London and Defence Science and Technology Laboratory. All human volunteers and dog were healthy. No signs of stress were observed from the dog during the experiments as the subject was already trained to wear a harness.

### 4.2. Fabrication of transducers and harness

We used silicone elastomer Ecoflex-30 (produced by Smooth-On, USA) in the fabrication of the transducers by mixing Part A and Part B in a 1:1 weight ratio. Once mixed, the mixture was degassed for 15 min in a vacuum chamber to remove any air bubbles before pouring into a mold and curing at room temperature.

#### Water-silicone Composite Diaphragm

The bottom part of the transducer was produced using a 3D printed PLA mold. The silicone piece was released from the mold after two hours of curing and filled with deionized water. Note that after two hours, the Ecoflex 30 could only partially cure as intended. We poured 5 gr of uncured Ecoflex 30 mixture on the surface of the water filled in the bottom silicone piece. The Ecoflex 30 spreads itself over water on its own, reaching the edges of the bottom silicone piece, and continues curing for another four hours into a composite structure. We have noticed that the silicone-water composite transducer loses small amounts of water over time (hence slightly decreased in size), potentially due to the porosity of silicone materials. Although this did not alter the experimental results over several months, this may be rectified by using liquids/gels with lower vapor pressure.

#### Air-silicone Composite Diaphragm

The bottom section of the air-silicone diaphragm was created by using the same procedure as described above. The top part of the diaphragm was created using a 3D printed mold that produces a silicone layer of 2 mm in thickness. Uncured Ecoflex 30 was poured into this new mold and a part-cured bottom section was placed in contact. The curing continued for another four hours.

#### All-silicone Diaphragm

Fabrication of the all-silicone piece was straight forward and involved replacing of water in the case of water-silicone composite, with uncured Ecoflex 30 that was poured into the part-cured Ecoflex 30 bottom piece. The entire piece was cured for another four hours to produce the final structure.

#### Assembly of transducers and harness

The final transducer structure was produced by placing a small printed circuit board (PCB) containing an electret microphone and a MAX9814 amplification chip on the bottom of the diaphragm and encasing the PCB once again with Ecoflex 30. Ecoflex 30 was also used to produce a harness with a 3D printed mold which was again co-cured with the transducer to create a monolithically integrated wearable device.

### 4.3 Mold production

The PLA molds used in the experiments were designed in SolidWorks and printed using a Raise3D N2 Plus 3D printer. The PLA filaments were purchased online from 3dgbire. The geometry of the molds was modeled after the Littman Classic II S.E. commercial stethoscope with a diameter of 45 mm.

### 4.4 Electronics & software

An ESP32-based microcontroller board (SparkFun ESP32 thing) was purchased online from SparkFun. The output of the audio amplifier MAX9814 was connected to the analog-to-digital converter (ADC) of ESP32. The audio signal was sampled at 8 kHz and samples were transmitted to a remote PC over Wi-Fi. In addition to remote transmission, we also designed an additional PCB containing an SD card module (**Figure S3**) using the EAGLE design software for off-line recording of digitized samples. The board was ordered from Elecrow Bazaar and surface mount components from Digikey. ECG samples were recorded using AD8232 - single lead heart rate monitor - also from Sparkfun. Schematic of the entire circuit is shown in **Figure S18**. We used MATLAB for creating a graphical interface and data processing. ESP32 was programmed using the PlatformIO integrated development environment with Arduino libraries.

### 4.5 Acquisition of Electrocardiogram

We have used SparkFun’s single-lead heart rate monitor (AD8232) sensor and microcontroller (ESP32) for the recording of the ECG signals. Simultaneous to ECG recording, we also recorded the heart sounds using the water-silicone composite transducer. The peaks of the ECG signal were detected using a modified version (see **Description SI-D2** for more information concerning our modifications and comparison of ECG and PCG recordings) of the algorithm described by Pan and Tompkins,^[39]^ Springer et. al,^[40]^ and Tosanguan et. al.^[41]^

### 4.6 Algorithmic detection of heart rate by PCG

We normalized the waveforms recorded with the water-silicone composite transducer between −1 and 1 and applied bandpass (passband: 20-150 Hz) filtering to eliminate any background noise. To localize S1 and S2 waveforms, we applied wavelet denoising, envelope detection and subsequent peak detection algorithms to find the peaks and calculate the distance between S1 signals to estimate the heart rate (for more information, please see **Description SI-D3**).

## Acknowledgements

Firat Güder would like to thank Imperial College Department of Bioengineering and the Institute for Security Science and Technology (funding under Champions Fund) and the Wellcome Trust (207687/Z/17/Z) for financial support. Yasin Cotur thanks the Turkish Ministry of Education. Matti Kaisti acknowledges Finnish Foundation for Technology Promotion. Michael Kasimatis acknowledges EPSRC DTP (Reference: 1846144). Firat Güder acknowledges Imperial College Centre for Plastic Electronics and EPSRC for Plastic Electronics Doctoral Training Centre (EP/G037515/1 and EP/L016702/1). Firat Güder also acknowledges Agri Futures Lab. We would like to thank Merve Cirisoglu Cotur for her help with the design of illustrations and videos. We would also like to thank Nina Cracknell, Stuart Cairns, Victoria Ratcliffe, and Hannah Robbins from Defense Science and Technology Laboratory (DSTL) for the fruitful discussions and their assistance for the experiments with dogs.

## Conflict of Interest

Authors declare no conflict of interest originating from this work.

## SYNOPSIS

**Flexible, stretchable, water-silicone composite** for acoustic transduction enables high performance real-time monitoring of vital signs over layers of clothing (or furry skin of animals) without the need for direct contact with the skin. Our method is based on an entirely new method of fabrication that involves encapsulation of water with silicones and exploits the ability of sound waves to travel through the body.

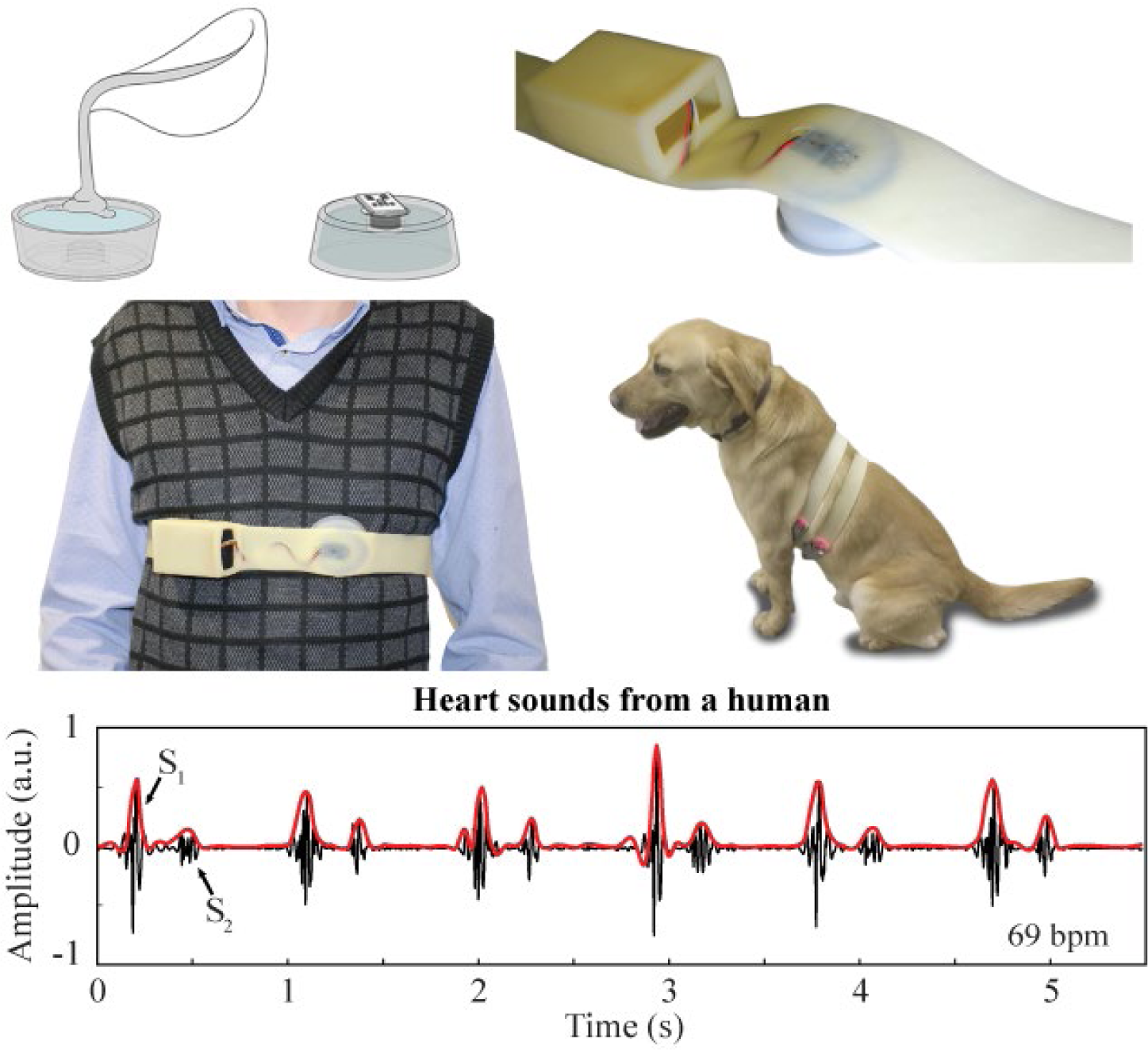

## References

[1] J. Andreu-Perez, D. R. Leff, H. M. D. Ip, G.-Z. Z. Yang, IEEE Trans. Biomed. Eng. 2015, 62, 2750.

[2] A. Pantelopoulos, N. G. Bourbakis, IEEE Trans. Syst. Man Cybern. Part C Appl. Rev. 2010, 40, 1.

[3] A. Kailas, C. C. Chong, F. Watanabe, IEEE Pulse 2010, 1, 57.

[4] A. M. Hussain, M. M. Hussain, Adv. Mater. 2016, 28, 4219.

[5] F. Güder, A. Ainla, J. Redston, B. Mosadegh, A. Glavan, T. J. Martin, G. M. Whitesides, Angew. Chemie - Int. Ed. 2016, 55, 5727.

[6] C. Dincer, R. Bruch, E. Costa-Rama, M. T. Fernández-Abedul, A. Merkoçi, A. Manz, G. A. Urban, F. Güder, Adv. Mater. 2019, 31.

[7] D. Maier, E. Laubender, A. Basavanna, S. Schumann, F. Güder, G. A. Urban, C. Dincer, ACS Sensors 2019, 4, 2945.

[8] C. Majidi, Soft Robot. 2014, 1, 5.

[9] X. Zhao, Q. Hua, R. Yu, Y. Zhang, C. Pan, Adv. Electron. Mater. 2015, 1.

[10] S. Sharma, A. El-Laboudi, M. Reddy, N. Jugnee, S. Sivasubramaniyam, S. M. El, P. Georgiou, D. Johnston, N. Oliver, A. E. G. G. Cass, M. El Sharkawy, P. Georgiou, D. Johnston, N. Oliver, A. E. G. G. Cass. Anal. Methods 2018, 10, 2088.

[11] A. J. Bandodkar, W. Jia, C. Yardimci, X. Wang, J. Ramirez, J. Wang, Anal. Chem. 2015, 87, 394.

[12] J. R. Windmiller, J. Wang, Electroanalysis 2013, 25, 29.

[13] J. H. So, A. S. Tayi, F. Güder, G. M. Whitesides, Adv. Funct. Mater. 2014, 24, 7197.

[14] O. Ranunkel, F. Güder, H. Arora, ACS Appl. Bio Mater. 2019, 2, 1490.

[15] X. Wang, Z. Liu, T. Zhang, Small 2017, 13, 1.

[16] E. Jo, K. Lewis, D. Directo, M. J. Y. Kim, B. A. Dolezal, J. Sport. Sci. Med. 2016, 15, 540.

[17] Z. Zhang, Z. Pi, B. Liu, IEEE Trans. Biomed. Eng. 2015, 62, 522.

[18] V. Marozas, A. Petrenas, S. Daukantas, A. Lukosevicius, J. Electrocardiol. 2011, 44, 189.

[19] R. Castrillón, J. J. Pérez, H. Andrade-Caicedo, Biomed. Eng. Online 2018, 17, 1.

[20] O. T. Inan, P. F. Migeotte, K. S. Park, M. Etemadi, K. Tavakolian, R. Casanella, J. Zanetti, J. Tank, I. Funtova, G. K. Prisk, M. Di Rienzo, IEEE J. Biomed. Heal. Informatics 2015, 19, 1414.

[21] M. Amjadi, K. U. Kyung, I. Park, M. Sitti, Adv. Funct. Mater. 2016, 26, 1678.

[22] S. Xu, Y. Zhang, L. Jia, K. E. Mathewson, K. I. Jang, J. Kim, H. Fu, X. Huang, P. Chava, R. Wang, S. Bhole, L. Wang, Y. J. Na, Y. Guan, M. Flavin, Z. Han, Y. Huang, J. A. Rogers, Science 2014, 344, 70.

[23] Y. Liu, J. J. S. Norton, R. Qazi, Z. Zou, K. R. Ammann, H. Liu, L. Yan, P. L. Tran, K. I. Jang, J. W. Lee, D. Zhang, K. A. Kilian, S. H. Jung, T. Bretl, J. Xiao, M. J. Slepian, Y. Huang, J. W. Jeong, J. A. Rogers, Sci. Adv. 2016, 2.

[24] Y. Shu, C. Li, Z. Wang, W. Mi, Y. Li, T. L. Ren, Sensors (Switzerland) 2015, 15, 3224.

[25] P. Chetlur Adithya, R. Sankar, W. A. Moreno, S. Hart. Biomed. Signal Process. Control 2017, 33, 289.

[26] V. Nivitha Varghees, K. I. Ramachandran, IEEE Sens. J. 2017, 17, 3861.

[27] B. S. Emmanuel, J. Med. Eng. Technol. 2012, 36, 303.

[28] N. Mohammadi-Koushki, H. Memarzadeh-Tehran, S. Goliaei, In Proceedings - Conference on Local Computer Networks, LCN; IEEE Computer Society, 2016; pp. 230–235.

[29] M. Nabih-Ali, E. S. A. El-Dahshan, A. S. Yahia, J. Med. Eng. Technol. 2017, 41, 553.

[30] Thinklabs Medical, Normal Heart Sound, https://www.thinklabs.com/heart-sounds, accessed: December 2019.

[31] H. Sakoe, S. Chiba, IEEE Trans. Acoust. 1978, 26, 43.

[32] M. Vlachos, M. Hadjieleftheriou, D. Gunopulos, E. Keogh, VLDB J. 2006, 15, 1.

[33] G. E. A. P. A. Batista, E. J. Keogh, O. M. Tataw, V. M. A. De Souza. Data Min. Knowl. Discov. 2014, 28, 634.

[34] P. Dickens, J. Smith, J. Wolfe, J. Acoust. Soc. Am. 2007, 121, 1471.

[35] R. A. Casarotto, J. C. Adamowski, F. Fallopa, F. Bacanelli, Arch. Phys. Med. Rehabil. 2004, 85, 162.

[36] O. A. Godin, J. Acoust. Soc. Am. 2008, 123, 1866.

[37] K. Vörös, I. Nolte, S. Hungerbühler, J. Reiczigel, J. Ehlers, G. Tater, R. Mischke, T. Zimmering, M. Schneider, Acta Vet. Hung. 2011, 59, 23.

[38] M. Kasimatis, E. Nunez-Bajo, M. Grell, Y. Cotur, G. Barandun, J.-S. Kim, F. Guder, ACS Appl. Mater. Interfaces 2019, acsami.9b17076.

[39] J. Pan, W. J. Tompkins, IEEE Trans. Biomed. Eng. 1985, 32, 230.

[40] D. B. Springer, T. Brennan, N. Ntusi, H. Y. Abdelrahman, L. J. Zühlke, B. M. Mayosi, L. Tarassenko, G. D. Clifford, J. Med. Eng. Technol. 2016, 40, 342.

[41] T. Tosanguan, PhD Thesis, Imperial College London, 2009.

